# Speech Processing Development in Infants: A Longitudinal ERP Study of Native and Nonnative Consonant Discrimination

**DOI:** 10.64898/2025.12.19.695368

**Authors:** Phillip M. Gilley, Daniel J. Tollin, Kristin M. Uhler

**Affiliations:** Department of Physical Medicine and Rehabilitation, University of Colorado Anschutz School of Medicine; Children’s Hospital Colorado, Aurora, CO USA; Department of Physiology & Biophysics, University of Colorado Anschutz Medical Campus; Institute of Cognitive Science, University of Colorado Boulder, Boulder, CO USA

**Keywords:** infant, ERP, mismatch response, perceptual attunement, speech perception, consonant discrimination, longitudinal

## Abstract

**Background:** Perceptual attunement theory predicts that during the first year of life, infants’ neural responses to native speech sounds should strengthen while responses to nonnative contrasts diminish. However, few longitudinal studies have directly tested these predictions using within-subject designs.

**Methods:** We recorded auditory event-related potentials (ERPs) from 84 typically developing infants at three time points (3, 6, and 12 months) using an oddball paradigm with native (/ba/-/da/) and nonnative (dental vs. retroflex /ʈa/) consonant contrasts. We examined developmental trajectories of the mismatch response (MMR) and obligatory components (P1, N1, P2) using linear mixed-effects models, with Session × Language interactions testing differential development for native versus nonnative contrasts. Center of mass analysis characterized spatial reorganization of ERP topography across development.

**Results:** Contrary to perceptual attunement predictions, most ERP measures showed similar developmental trajectories for native and nonnative consonants. Of 14 interaction tests, only N1 latency for deviant waveforms showed a significant Session × Language interaction (p = 0.046, ηp² = 0.025). Significant main effects of Session on multiple measures confirmed ongoing auditory maturation. Center of mass analysis revealed systematic spatial reorganization: P1 showed posterior migration and leftward shifts (0.078 units displacement), while N1 and MMR exhibited anterior shifts, with nonnative consonants showing larger spatial reorganization for N1 (0.099 units vs. 0.058 units for native).

**Conclusions:** Robust neural signatures of perceptual attunement for consonants may be more subtle, later-emerging, or stimulus-specific than commonly assumed. The dissociation between stable amplitude and migrating generators suggests that spatial location may be as important as response magnitude for characterizing auditory maturation during early infancy. Practically, equivalent neural responses to native and nonnative consonant contrasts should not be interpreted as evidence of atypical development; rather, indices of general auditory maturation may be more reliable markers of healthy development during the first year.

## 1. Introduction

Human infants are born with a broad capacity to discriminate speech sounds as documented by both neurophysiological and perceptual studies. Over the course of the first year of life, discrimination of speech sounds is shaped by the child’s home language(s). Research has shown that these perceptual changes are first observed for vowels (Kuhl et al., 1992; Strange & Jenkins, 1978), followed by enhanced abilities to differentiate native consonants a few months later (Werker & Tees, 1984). This transition is commonly referred to as perceptual attunement (or perceptual narrowing), in which maturational changes in auditory cortical circuitry interact with experience-dependent plasticity driven by the statistics of the infant’s language environment (Reh et al., 2020). Converging methods support this account: EEG studies demonstrate age- and experience-linked reorganization of neural responses to phonetic changes (Peña et al., 2012; Petitto et al., 2013; Werwach et al., 2022), neuroimaging points to early specialization for native speech and experience-related changes in functional responses (Cabrera & Gervain, 2020; Zhao & Kuhl, 2022), and behavioral paradigms reveal language-specific tuning for native vowels by ∼6 months with strengthening sensitivity to many native consonant contrasts by ∼9-12 months (Kuhl et al., 1992; Strange & Jenkins, 1978; Ortiz-Mantilla et al., 2016; Werker & Tees, 1984; Reh et al., 2021). An additional important hallmark of this process is reduced discrimination of non-native speech as native categories sharpen (Kuhl et al., 1992; Werker & Tees, 1984).

Importantly, variability in attunement trajectories has clinical consequences. Infants who do not show strengthening discrimination for native contrasts during early development tend to have poorer subsequent language outcomes, highlighting the first year as a sensitive period for auditory development and later language learning (Tsao et al., 2004; Rivera-Gaxiola et al., 2005; Conboy & Kuhl, 2008; Kuhl, 2010). Yet, despite broad agreement on the existence of a sensitive period, reported timelines differ across methods and paradigms, leaving the precise course of development an open question (Peña et al., 2012; Petitto et al., 2013; Reh et al., 2020). These methodological discrepancies motivate a closer integration of behavioral and neurophysiological measures to delineate what changes when, and why.

A complementary thread in developmental cognitive neuroscience concerns how auditory and language functions become lateralized across the two cerebral hemispheres. Research using functional magnetic Magnetic Resonance Imaging, functional Near-Infrared Spectroscopy (fNIRS)/optical topography, and combined fNIRS-EEG has revealed that hemispheric specialization for speech is evident already at birth: both hemispheres respond to speech, but the right hemisphere preferentially encodes spectral and slow temporal/prosodic aspects, whereas the left hemisphere is relatively more sensitive to faster temporal modulations that support segmental and phonological processing (Zatorre & Belin, 2001; Dehaene-Lambertz et al., 2002; Peña et al., 2003; Homae et al., 2006, 2007; Arimitsu et al., 2011; Friederici, 2011; Minagawa-Kawai et al., 2011; Bortfeld et al., 2009; Cabrera & Gervain, 2020; Perani et al., 2011). Across infancy and early childhood, functional imaging further suggests a shift from broad, bilateral recruitment toward increasingly localized - and increasingly left-lateralized - responses as experience with native speech sounds accumulates (Dehaene-Lambertz et al., 2002; Holland et al., 2007). Thus, while attunement describes what is changing in the representational landscape, lateralization helps specify where in the brain those changes consolidate.

The ideas of attunement and hemispheric specialization converge on a coherent developmental trajectory. In the first days of life, phonemic deviants predominately engage the left hemisphere and prosodic deviants the right, revealing lateralized specialization prior to extensive language exposure (Arimitsu et al., 2011). By approximately three months, the left hemisphere already shows adult-like selectivity for structured speech, while the right hemisphere robustly encodes spectral/slow temporal information such as intonation (Dehaene-Lambertz et al., 2002; Homae et al., 2006). By ∼4-6 months, language-specific tuning becomes detectable and relatively left-hemisphere lateralized for native compared with non-native speech when contrasted with non-speech controls, and this lateralization remains robust in naturalistic awake paradigms (Bortfeld et al., 2009; Minagawa-Kawai et al., 2011). Together, these findings support a time course in which a bilateral fronto-temporal cortical system present at birth exhibits a right-hemisphere advantage for prosody and a left-hemisphere bias for faster modulations and lawful linguistic structure; experience with native speech then sharpens these biases while white-matter and functional connectivity mature to consolidate a left-dominant network that nonetheless preserves right-hemisphere specialization for prosody (Perani et al., 2011; Minagawa-Kawai et al., 2011; Cabrera & Gervain, 2020).

Framed against this backdrop, debates about the pace and endpoint of auditory maturation become increasingly important to consider as we strive towards a measure that could inform future interventions. With these goals in mind, we explored two divergent auditory development models: stability model and incremental model. The stability model proposes that core auditory development is largely complete by middle childhood, pointing to near-adult detection and frequency-discrimination abilities by ∼6 years and functional maturity of Heschl’s gyrus by ∼7 years (Maxon & Hochberg, 1982; Moore, 2002; Sharma et al., 1997). In contrast, the incremental model posits continued improvements into adolescence, consistent with protracted gains in higher-order listening (e.g., speech-in-noise) and prolonged myelination/synaptic pruning in secondary auditory cortex (Ponton et al., 2000; Moore & Linthicum, 2007). An unresolved issue critical for interpreting both attunement and hemispheric lateralization is whether age-related changes primarily reflect bottom-up physiological maturation or increasing reliance on top-down strategies and attentional control or some combination of both (Bishop et al., 2011).

To address these questions speech perception must be assessed during the first two years of life, a developmental window that is not only theoretically informative but also clinically significant for early identification of risk for perception development, informing habilitation plans and evaluation of intervention efficacy. To study discrimination mechanisms and chart their developmental trajectories, objective tools such as electrophysiology and neuroimaging are essential. Among these, auditory event-related potentials (ERPs) are especially well suited: they are noninvasive, temporally precise, and administered in sleeping or awake infants. Developmental ERP studies document age-related changes in obligatory components such as P1 and the T-complex thought to index partially parallel auditory pathways with a large, broadly distributed P1 in infants/children and prominent temporal-site T-complex generators in childhood that diminish with age (Sharma et al., 1997; Ponton et al., 2000; Bishop et al., 2011). ERP morphology is sensitive to stimulus class, rate, intensity, and developmental stage, and trajectories can be step-like or gradual, underscoring the value of longitudinal and cross-sectional designs alike.

The mismatch responses/negativity (MMR/MMN) is an ERP elicited in auditory oddball paradigms which offers a powerful approach to probing early speech discrimination and its reorganization with experience such as a child’s spoken home language. Oddball designs compare neural responses to frequently presented standard tokens and infrequent deviant tokens that differ along controlled phonetic dimensions. Crucially for infant research, speech-evoked MMR/MMN can be recorded during natural sleep, require no overt behavior, and can be configured to test both native and non-native contrasts in a single session by manipulating the identity and magnitude of deviants. Prior work shows that infant MMRs capture core features of perceptual attunement: responses to native deviants generally strengthen and become more adult-like in the latter half of the first year, whereas responses to non-native deviants diminish or exhibit polarity patterns consistent with less salient or “harder” changes. Moreover, individual differences in MMR/MMN to native contrasts between ∼7-11 months predict later language outcomes, including vocabulary growth, suggesting clinically useful prognostic value (Conboy & Kuhl, 2008).

These past studies, including our own, provide the building blocks to leverage ERPs in an auditory oddball paradigm to connect three strands that are often examined separately: perceptual attunement, the maturation of hemispheric specialization, and objective electrophysiologic markers of auditory development. By quantifying early speech-evoked ERPs to both standards and deviants across native and non-native consonant contrasts in infants aged 3-12 months, we aim to dissociate contributions of auditory system maturation from experience with specific speech cues.

## 2. Methods

### 2.1. Participants

Participants were 83 typically developing infants enrolled in an accelerated longitudinal study examining speech perception development across the first year of life. Data were collected at three time points: T1 (3 months of age), T2 (6 months), and T3 (12 months). At T1, 59 infants contributed data. At T2, 72 infants participated. At T3, 46 infants completed testing.

The final analyzed dataset included ERP recordings distributed across three sessions and both consonant stimulus conditions. Sample sizes varied by condition due to data quality filtering and the number of children who had completed each session.

Per parent report, all infants passed newborn hearing screening assessed by auditory brainstem response and had no reported history of neurologic disorders or hearing loss. Prior to participation, parents received written and oral information describing the study aims and procedures and provided written informed consent. All procedures conformed to the ethical standards of the Declaration of Helsinki (1964) and subsequent revisions, and the protocol was approved by the Colorado Multiple Institutional Review Board (COMIRB, protocol number: 22-2374).

### 2.2. Procedures

Testing sessions lasted approximately 45 minutes, although three hours were scheduled for each participant to accommodate infant needs. Middle ear status was evaluated via tympanometry using age-appropriate probe tone and otoacoustic emissions. Infants were either placed in a rocker or held by a parent in a quiet, dimly lit room to facilitate or maintain sleep. The rocker’s motion remained inactive during EEG recordings. At the third time point (T3), if the infant did not fall asleep, engagement was maintained with a silent video or toys. Previous research has demonstrated reliable elicitation of the mismatch response (MMR) across a range of infant sleep states (Martynova et al., 2003; Sambeth et al., 2009; Uhler et al., 2018).

Electrophysiological recordings were obtained from 11 Ag/AgCl electrodes positioned according to the International 10-20 system (F5, Fz, F6, C5, Cz, C6, P5, Pz, P6, M1, & M2) and referenced to the nasion (Nz). A bipolar channel was placed at the lateral canthus of the right eye and referenced to the superior orbit to monitor ocular activity and arousal. When infants were held by a parent, the ground electrode was placed on the parent’s forearm.

Data were recorded using either a SynAmps2 EEG amplifier (Compumedics Neuroscan, Charlotte, NC, USA) or a MindfulMobile EEG amplifier (Neuracle Neuroscience, Beijing, China). Continuous EEG was acquired at a sampling rate of 10000 Hz (SynAmps2) or 4000 Hz (MindfulMobile) and band-pass filtered from DC to 4000 Hz during each experimental block. The high sampling rate and band-pass filter was chosen to accommodate a secondary analysis for a different study.

Parents received $100 compensation per completed visit, and parking costs were reimbursed by the University. All procedures conformed to the ethical standards of the Declaration of Helsinki (1964) and subsequent revisions, and the protocol was approved by the University of Colorado Institutional Review Board (protocol number: 22-2374).

### 2.3. Stimuli and Experimental Design

Cortical auditory evoked potentials (CAEPs) were recorded during passive listening to speech sound contrasts using an auditory oddball paradigm. Two consonant stimulus contrasts were presented in separate blocks: (1) Native Consonant (/ba/ vs. /da/), and (2) Nonnative Consonant (dental vs. retroflex /ʈa/). Within each block, standard stimuli were presented with 85% probability and deviant stimuli with 15% probability, with the constraint that deviant stimuli never appeared in succession. Stimuli were 400 ms in duration, presented in the sound field at 70 dBA with an inter-stimulus interval of 1200 ms. Approximately 600 trials (∼510 standard, ∼90 deviant) were collected per block.

### 2.4. EEG Recording and Preprocessing

Continuous EEG data were first subjected to a series of quality control and preprocessing steps to ensure signal integrity prior to epoching. Data were downsampled to 1000 Hz to standardize the temporal resolution across the dataset. The continuous EEG was then examined for bad channels using automated detection algorithms that identified electrodes with abnormally high impedance, excessive noise, or poor contact with the scalp. Following automated channel identification, the continuous data were visually inspected to ensure that the recorded waveforms exhibited typical characteristics and to identify any periods of recording contaminated by equipment malfunction or extreme non-neural artifacts that would compromise subsequent analyses.

Spectral filtering was applied to the continuous EEG prior to epoching using a zero-phase 4th-order Butterworth bandpass filter with cutoff frequencies of 1 Hz (high-pass) and 18 Hz (low-pass). This filter specification was selected based on our previous work demonstrating optimal signal-to-noise characteristics for infant cortical auditory evoked potentials within this frequency range (Gilley et al., 2017; Uhler et al., 2018). The decision to filter the continuous data rather than individual epochs was made to avoid edge artifacts that can arise when filters are applied to short time windows, particularly given the finite impulse response characteristics of the Butterworth filter.

Following spectral filtering, the continuous EEG was segmented into epochs from -200 to +799 ms relative to stimulus onset. Artifact rejection was then performed on the epoched data using a combination of automated and manual procedures. Automated rejection criteria included threshold-based detection of extreme voltages (±150 μV), step-like artifacts (>50 μV voltage difference between consecutive samples), and abnormally distributed data (kurtosis >5 SD from the mean). Additional manual inspection was conducted to identify epochs contaminated by movement artifacts, muscle activity, or other sources of non-neural electrical activity.

Following artifact rejection, epoched data were downsampled to 250 Hz to reduce computational load while maintaining adequate temporal resolution for ERP component identification. Baseline correction was applied by subtracting the mean voltage during the pre-stimulus interval (-200 to 0 ms) from each time point in the epoch. To further reduce spatially uncorrelated noise, spatial principal component analysis (PCA) was applied across channels. This procedure involved z-score normalization of the data across channels, followed by eigendecomposition of the channel covariance matrix. Components accounting for 95% of the cumulative variance were retained and projected back to the electrode space, effectively removing noise sources such as isolated electrode artifacts while preserving the primary spatial patterns of the evoked response. Following PCA reconstruction and de-normalization, baseline correction was re-applied to ensure accurate voltage referencing.

#### 2.4.1. Trial Matching

For all remaining analyses, standard trials were selected to match the number of deviant trials within each experimental condition to ensure balanced comparisons. The default matching procedure employed a “preceding standard” strategy where, for each deviant trial, the immediately preceding standard trial was selected. This approach maintains temporal proximity between matched trial pairs and controls for potential time-on-task effects such as changes in arousal or sleep state. An alternative random permutation method, in which standard trials were randomly sampled without replacement to match deviant trial counts, was also available to verify that the matching strategy did not systematically bias the results.

#### 2.4.2. ERP Waveform Estimation and Statistical Analysis

For each experimental condition defined by consonant type (native vs. nonnative) and session (T1, T2, T3), mean ERP waveforms and 95% confidence intervals (CI) were estimated using bootstrap resampling procedures. Bootstrap samples were drawn with replacement from available trials across 10,007 iterations, and the condition mean was computed as the average across bootstrap iterations. CI (95%) were derived from the 2.5th and 97.5th percentiles of the bootstrap distribution at each channel and time point. Standard, Deviant, and Mismatch Response (MMR; computed as Deviant minus Standard) waveforms were generated using these bootstrap procedures for each participant, condition, and session. Figure 1 displays the Standard and Deviant ERP waveforms and CI for each condition and session. Figure 2 displays the MMR waveforms and CI for each condition and session.

**Figure 1.**
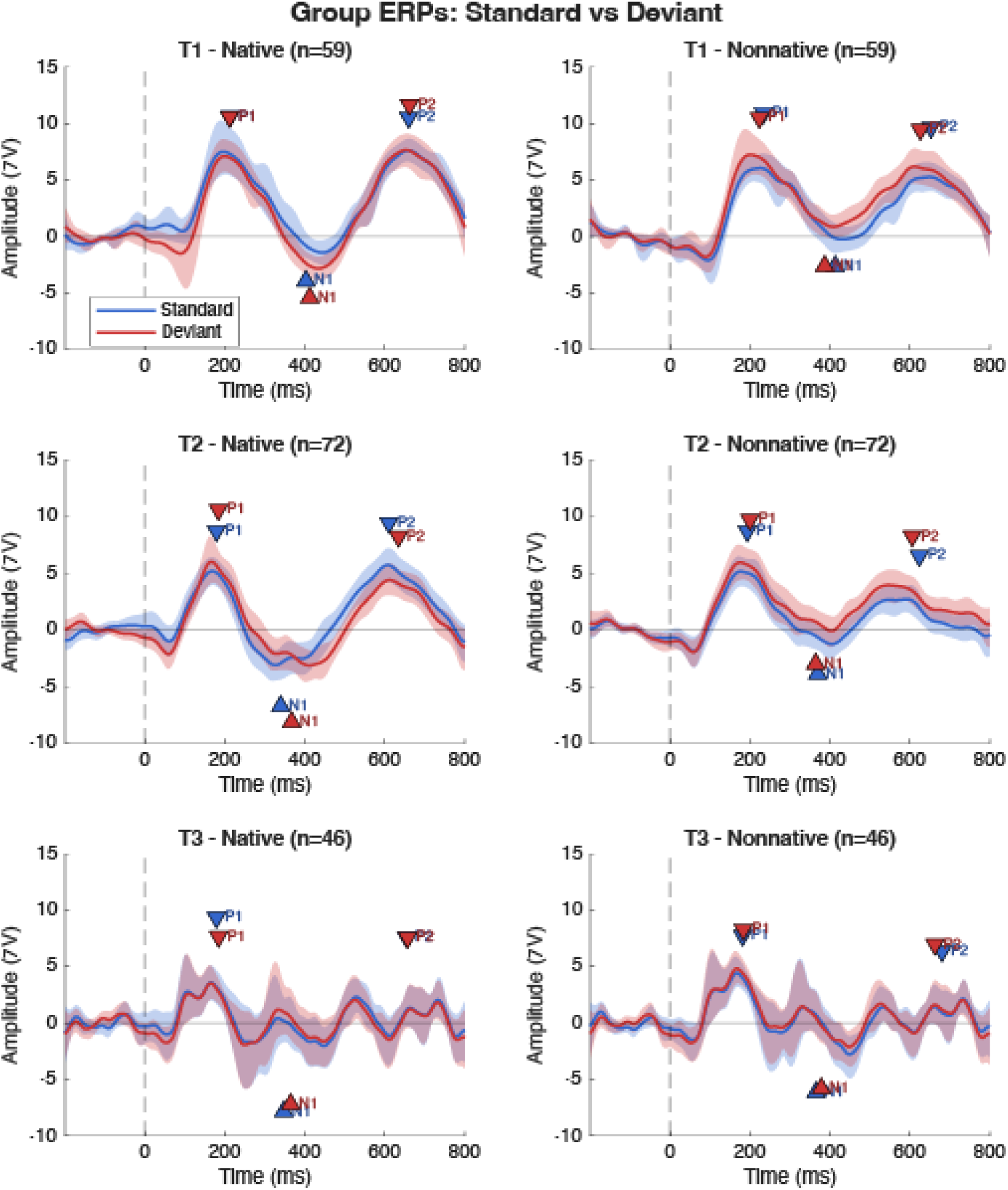
Group ERPs for Standard and Deviant responses for each Session x Language interaction. Each panel displays the Standard mean in blue and the Deviant mean in red. Shaded regions represent the 95% confidence intervals. Peak markers for the P1, N1, and P2 peaks represent the average latency and amplitude of the picked peak locations.

**Figure 2.**
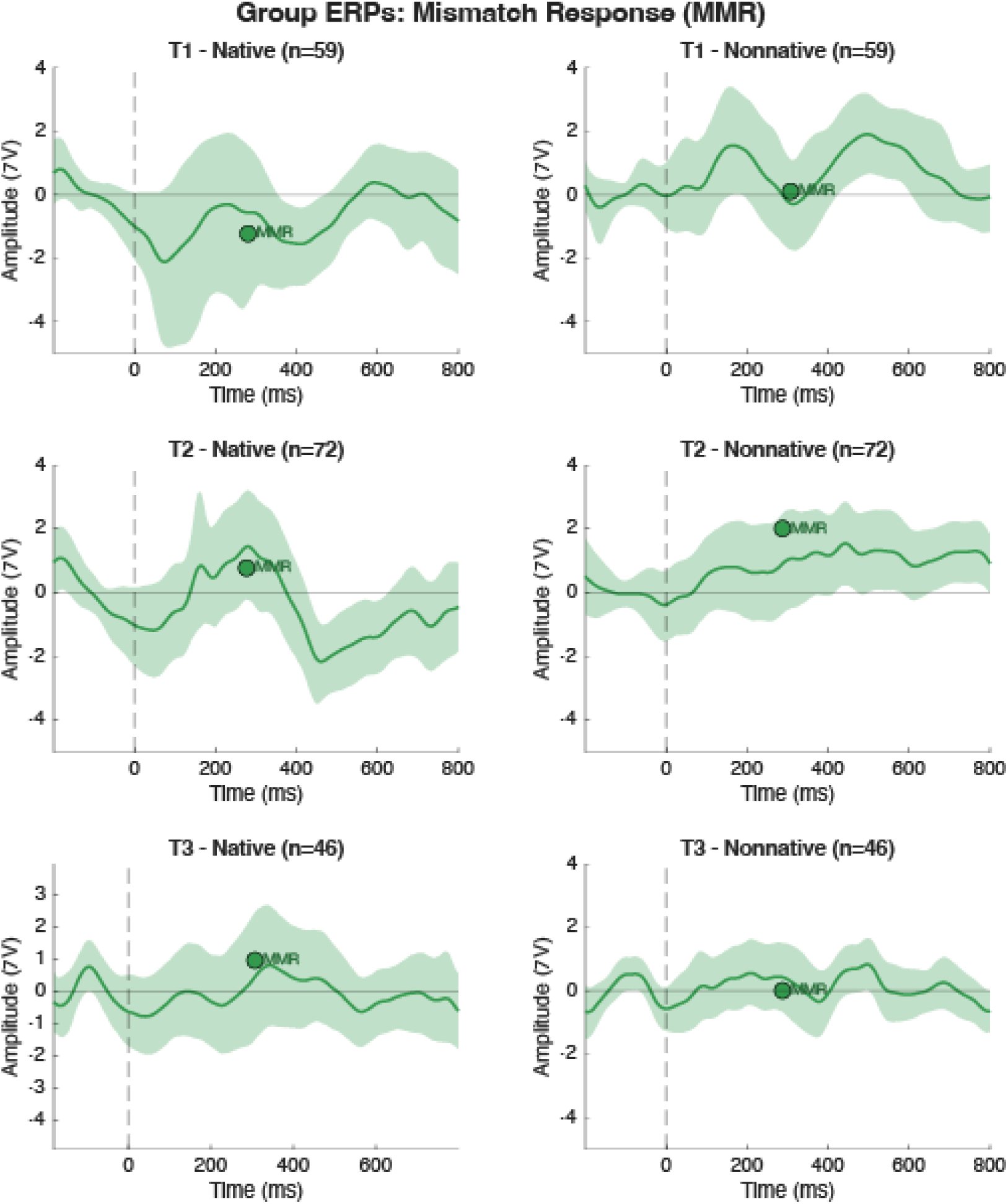
Group ERPs for the mismatch response (MMR) for each Session x Language interaction. Each panel displays the mean MMR in green. Shaded regions represent the 95% confidence intervals. Peak markers for the MMR represent the average latency and amplitide of the picked MMR peak locations.

Statistical hypothesis testing of the MMR was conducted using two complementary approaches: Bayesian analysis and frequentist independent-samples t-tests. Bayesian analysis computed Bayes Factors (BF₁₀) at each time point, quantifying the strength of evidence for a difference between deviant and standard responses relative to the null hypothesis of no difference. BF₁₀ > 3 was interpreted as substantial evidence for an MMR, and BF₁₀ > 10 as strong evidence (Kass & Raftery, 1995). Independent-samples t-tests comparing standard and deviant trial distributions at each time point were corrected for multiple comparisons using the False Discovery Rate (FDR) procedure (Benjamini & Hochberg, 1995) to control the expected proportion of false discoveries across the full time series.

### 2.5. ERP Component Identification

ERP component peaks (P1, N1, P2 for standard and deviant waveforms; peak MMR for mismatch responses) were identified from a vertex montage computed as the average of Fz and Cz channels re-referenced to linked mastoids (M1/M2). This vertex montage approach was selected based on our previous work demonstrating reliable detection of infant cortical auditory evoked potentials at fronto-central electrode sites (Gilley et al., 2017). The linked mastoid reference provides a stable reference that minimizes contamination from non-neural sources while preserving the characteristic morphology of infant auditory ERPs. Averaging the Fz and Cz channels improves signal-to-noise ratio by combining activity from two sites that typically show maximal amplitude for the P1-N1-P2 complex in infants.

Peaks were identified from the vertex waveform using component-specific search windows derived from the grand average of all waveforms for all participants: P1 (100-300 ms), N1 (200-450 ms), P2 (500-800 ms), and peak MMR (150-450 ms). Local maxima (for positive components) and minima (for negative components) were identified within each search window, with a minimum peak distance constraint of 80 ms enforced to prevent double-counting of a single component. Peak prominence and signal-to-noise ratio were used to select the most robust deflection when multiple candidate peaks were present. Peak amplitudes and latencies extracted from the vertex montage were subsequently treated as the response variables for statistical analysis.

### 2.6. Spatial Analysis of Scalp Topography

To examine developmental changes in the scalp distribution of ERP components, we computed the amplitude-weighted center of mass for each peak at each session using three-dimensional electrode coordinates normalized to a unit sphere. The center of mass (CoM) was calculated as the weighted average of channel coordinates, with weights proportional to the absolute peak amplitude averaged across subjects at each channel:

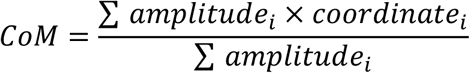

where *i* indexes the 11 recording channels. Spatial coordinates were defined relative to a standard three-dimensional coordinate system (X-axis: left/right, Y-axis: posterior/anterior, Z-axis: inferior/superior) on a normalized unit sphere with radius 1. This normalization allowed spatial shifts to be quantified in standardized units that are comparable across sessions despite age-related changes in head size and electrode placement.

Developmental shifts in scalp topography were quantified as the Euclidean distance between the center of mass at T1 (baseline) and subsequent sessions (T2, T3), with directional components (ΔX, ΔY, ΔZ) computed separately to characterize lateral, anterior-posterior, and inferior-superior shifts. Shifts were characterized in normalized units as minimal (< 0.02), small (0.02–0.05), moderate (0.05–0.10), or large (≥ 0.10) based on the total displacement magnitude.

It is important to note that these spatial measures characterize changes in the surface distribution of voltage patterns recorded at the scalp rather than directly localizing underlying neural generators. Given the limited spatial sampling afforded by 11 normalized electrode positions, the present analysis cannot support strong inferences about specific cortical sources or their reorganization. Nevertheless, systematic shifts in scalp topography across development may reflect changes in the relative contributions of different cortical areas, alterations in the orientation or strength of underlying dipole configurations (e.g., T-complex), or developmental changes in head geometry that affect volume conduction to the scalp. The topographic analysis thus provides a complementary perspective on developmental changes in auditory processing, possibly showing that functional reorganization indexed by amplitude and latency measures is accompanied by reliable shifts in the spatial distribution of scalp-recorded activity. Figure 3 displays the mean scalp topographies at for each peak (P1, N1, P2, and MMR) at each session for responses to native speech sounds.

**Figure 3.**
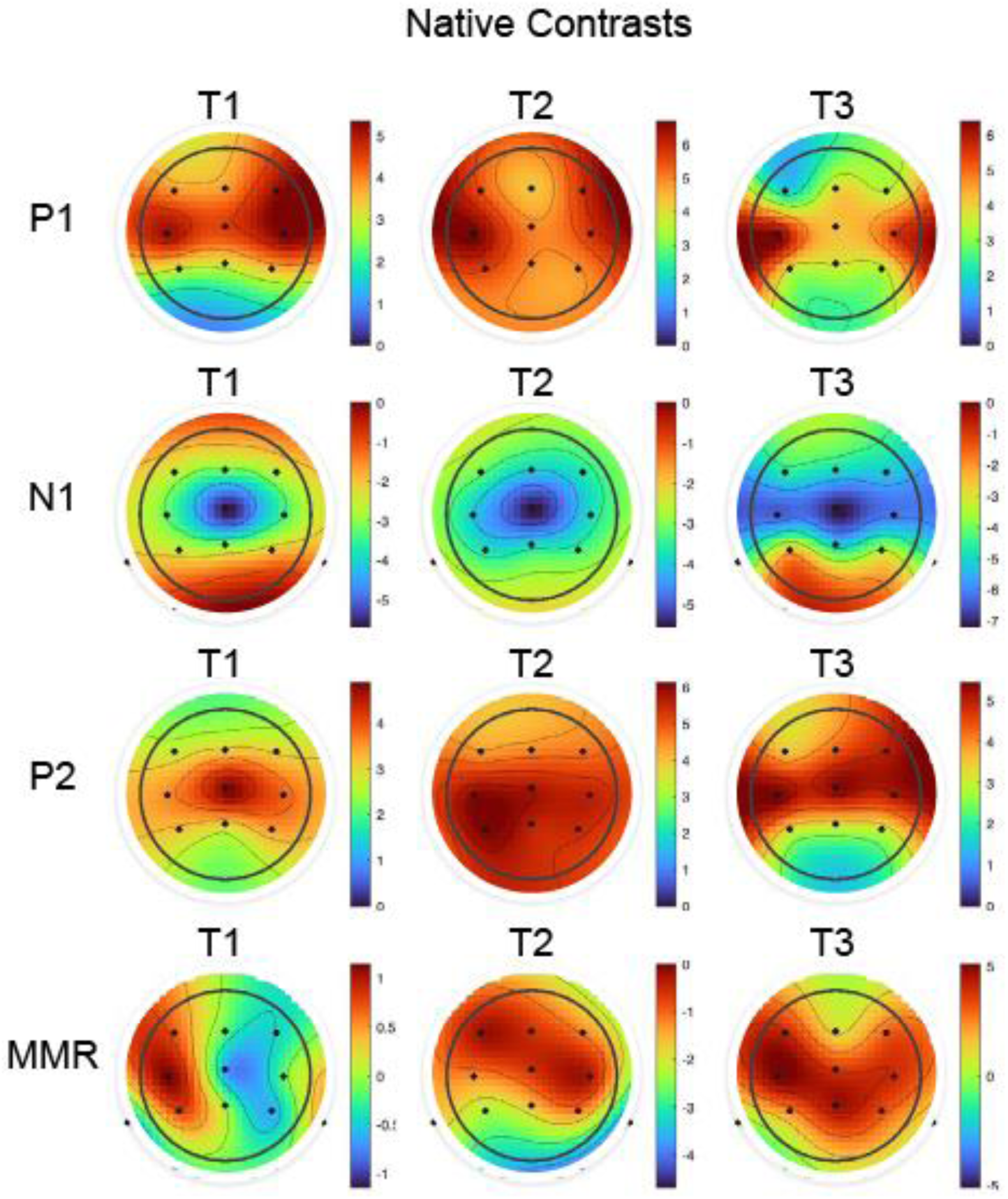
Group mean topographic scalp distributions of the Native contrasts for each peak (P1, N1, P2, and MMR; rows) and for each Session (T1, T2, and T3; columns).

**Figure 4.**
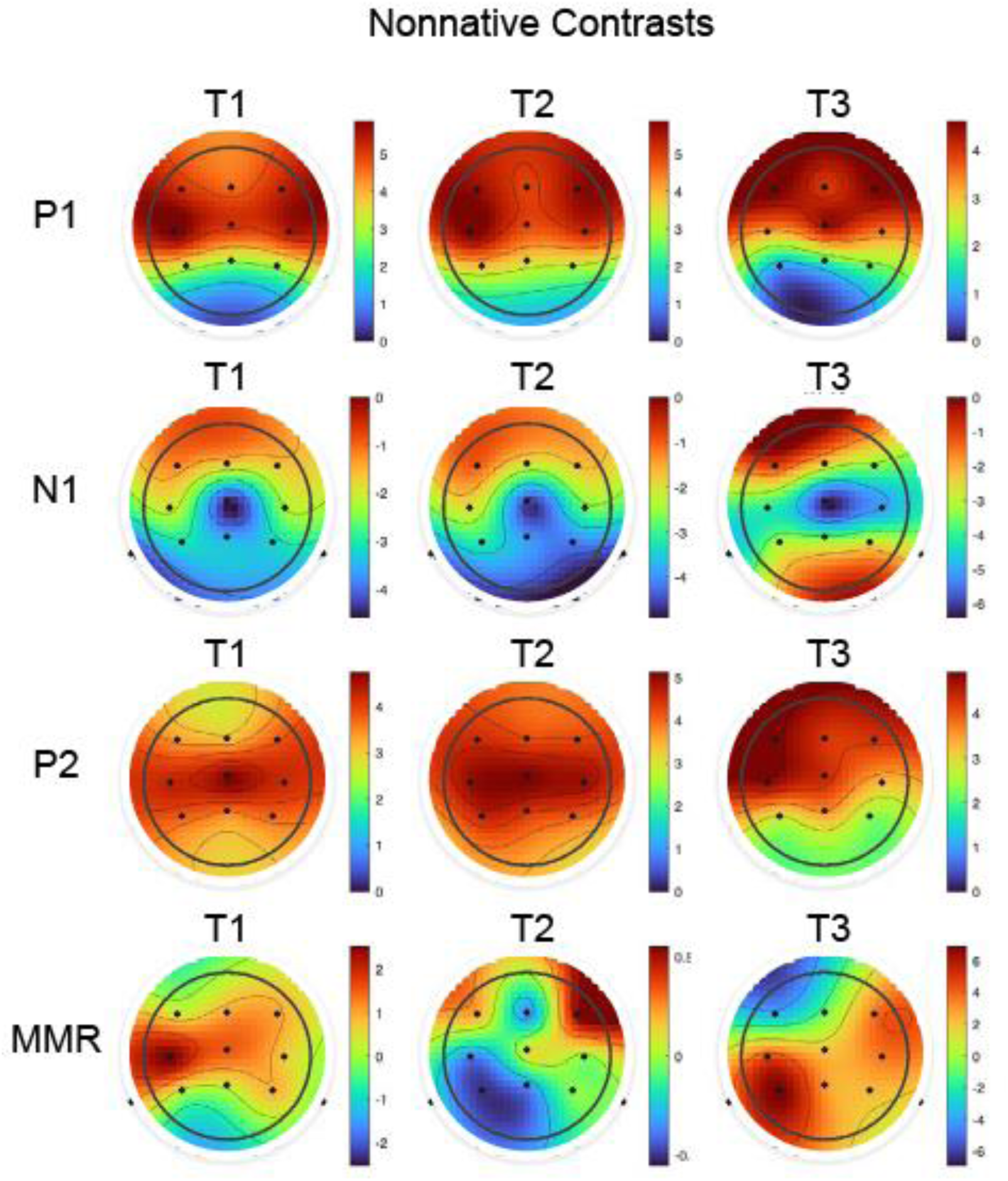
Group mean topographic scalp distributions of the Nonnative contrasts for each peak (P1, N1, P2, and MMR; rows) and for each Session (T1, T2, and T3; columns).

### 2.7. Statistical Analysis

Developmental changes in ERP component latencies and amplitudes were assessed using linear mixed-effects (LME) models implemented in MATLAB (R2025b, Statistics and Machine Learning Toolbox). Mixed-effects models accommodate the unbalanced longitudinal design, include all available data (maximizing statistical power), and account for repeated measures within participants.

#### 2.6.1. Primary Analysis: Developmental Trajectory Comparison

The primary research question was whether native and nonnative consonant contrasts show different developmental trajectories across the first year of life. To address this question directly, we specified LME models with Session, Language (Native vs. Nonnative), and their interaction as fixed effects:

##### MMR Analysis

MMR_Amplitude ∼ Session × Language + (1 | SubjectID)

MMR_Latency ∼ Session × Language + (1 | SubjectID)

##### ERP Peak Analysis (P1, N1, P2)

For each peak, separate models were fit for Standard and Deviant waveforms:

Amplitude ∼ Session × Language + (1 | SubjectID)

Latency ∼ Session × Language + (1 | SubjectID)

The critical test of differential development is the Session × Language interaction: a significant interaction indicates that native and nonnative contrasts follow different developmental trajectories, whereas a non-significant interaction suggests similar developmental patterns regardless of linguistic relevance.

F-statistics, degrees of freedom, and p-values were computed using ANOVA with Satterthwaite approximation for denominator degrees of freedom. Partial eta-squared (ηp²) effect sizes were computed for each effect.

#### 2.6.2. Bayes Factor Analysis

To complement the frequentist analysis, Bayesian analysis was conducted on individual subject data. For each subject, session, and language condition, we computed the maximum Bayes Factor (BF₁₀) across the MMR time window, representing the peak evidence for neural discrimination. These values were log-transformed and analyzed using LME models with the same Session × Language factorial structure to test whether evidence strength showed differential developmental trajectories.

#### 2.6.3. Effect Size Interpretation

Partial η² values were interpreted as negligible (< 0.01), small (0.01-0.06), medium (0.06-0.14), or large (≥ 0.14).

#### 2.6.4. Software Implementation

All analyses were conducted in MATLAB R2025b (The MathWorks, Natick, MA) using the Statistics and Machine Learning Toolbox. Linear mixed-effects models were fit using the fitlme function, ANOVA F-tests were extracted using the anova function with Satterthwaite degrees of freedom. The significance threshold was set at α = 0.05 for all statistical tests.

## 3. Results

### 3.1. Sample Description

The final analyzed sample included 83 unique subjects contributing 178 total sessions across the three time points. Session-specific sample sizes were: T1 (n = 59), T2 (n = 72), and T3 (n = 46). Each subject contributed data to both Native and Nonnative conditions within each session attended.

### 3.2. MMR Analysis

#### 3.2.1. MMR Amplitude (Primary Outcome)

The primary analysis examined whether MMR amplitude showed differential development between native and nonnative consonant contrasts. The LME model included 356 observations from 84 subjects.

The Session × Language interaction was not significant (F(2, 271.8) = 0.41, p = 0.667, ηp² = 0.003), indicating that native and nonnative contrasts showed similar developmental trajectories for MMR amplitude. Neither the main effect of Session (F(2, 308.8) = 0.73, p = 0.480, ηp² = 0.005) nor the main effect of Language (F(1, 271.8) = 0.51, p = 0.476, ηp² = 0.002) reached significance.

Descriptive statistics revealed that MMR amplitude remained relatively stable across development for both conditions (Table 1). For native contrasts, mean amplitude was -1.23 μV (SD = 12.93) at T1, 0.77 μV (SD = 12.65) at T2, and 0.95 μV (SD = 8.54) at T3. For nonnative contrasts, mean amplitude was 0.11 μV (SD = 10.93) at T1, 2.01 μV (SD = 9.18) at T2, and 0.00 μV (SD = 5.94) at T3.

**Table 1.** MMR Amplitude Descriptive Statistics.

| Session | Language | Mean (μV) | SD | N |
| --- | --- | --- | --- | --- |
| T1 | Native | -1.23 | 12.93 | 59 |
| T1 | Nonnative | 0.11 | 10.93 | 59 |
| T2 | Native | 0.77 | 12.65 | 72 |
| T2 | Nonnative | 2.01 | 9.18 | 72 |
| T3 | Native | 0.95 | 8.54 | 46 |
| T3 | Nonnative | 0.00 | 5.94 | 46 |

#### 3.2.2. MMR Latency (Secondary Outcome)

MMR latency was examined as a secondary outcome indexing the timing of neural discrimination. The Session × Language interaction was not significant (F(2, 356.0) = 1.47, p = 0.231, ηp² = 0.008), indicating similar developmental trajectories. Neither the main effect of Session (F(2, 356.0) = 1.46, p = 0.233, ηp² = 0.008) nor Language (F(1, 356.0) = 2.24, p = 0.136, ηp² = 0.006) was significant.

Descriptive statistics showed stable MMR latencies across development (Table 2). For native contrasts, mean latency was 280 ms (SD = 109) at T1, 277 ms (SD = 95) at T2, and 307 ms (SD = 94) at T3. For nonnative contrasts, mean latency was 307 ms (SD = 105) at T1, 287 ms (SD = 98) at T2, and 287 ms (SD = 90) at T3.

**Table 2.**
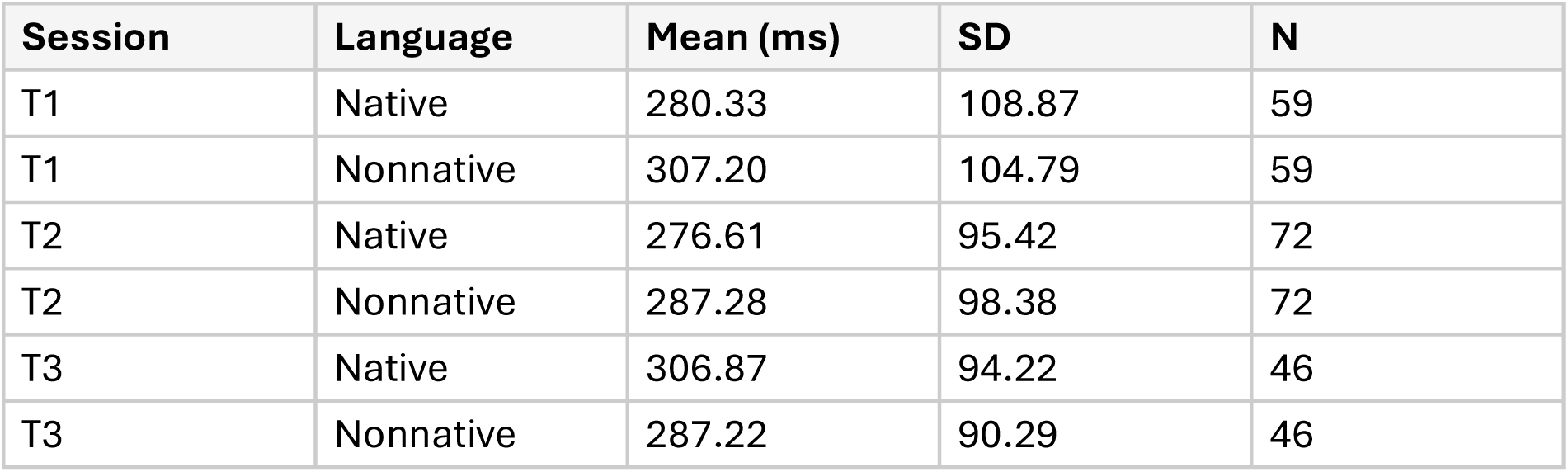
MMR Latency Descriptive Statistics.

#### 3.2.3. MMR Spatial Characteristics (Center of Mass)

Center of mass analysis revealed systematic anterior shifts in the topographic distribution of the MMR response from T1 to T3 for both language conditions. For native consonants, the total 3D displacement was 0.061 units, with the anterior-posterior component accounting for the majority of this shift (0.059 units anterior). The trajectory showed an initial small posterior shift from T1 to T2 (0.018 units) followed by a larger anterior shift from T2 to T3, resulting in net anterior displacement of the topographic distribution over the full developmental interval. For nonnative consonants, the MMR topographic distribution showed substantial anterior shifts from T1 to T3 (0.065 units total, with 0.063 units in the anterior-posterior dimension), closely paralleling the pattern observed for native consonants. The magnitude and direction of this topographic shift were remarkably similar across the two consonant conditions.

### 3.3. ERP Peak Analysis

ERP peaks (P1, N1, P2) were analyzed separately for Standard and Deviant waveforms to examine whether obligatory auditory responses showed differential developmental trajectories between native and nonnative conditions.

#### 3.3.1. P1 Component

##### P1 Latency

The Session × Language interaction was not significant for either Deviant (F(2, 249.0) = 1.23, p = 0.295, ηp² = 0.010) or Standard waveforms (F(2, 257.0) = 0.89, p = 0.413, ηp² = 0.007). A significant main effect of Session was observed for both waveforms (Deviant: F(2, 277.8) = 8.24, p < 0.001, ηp² = 0.056; Standard: F(2, 288.8) = 9.09, p < 0.001, ηp² = 0.059), indicating general maturational changes in P1 timing that did not differ by language condition. For Standard waveforms, a significant main effect of Language emerged (F(1, 254.8) = 5.77, p = 0.017, ηp² = 0.022), with native contrasts showing shorter latencies overall.

##### P1 Amplitude

No significant Session × Language interactions were observed for either Deviant (F(2, 265.6) = 0.18, p = 0.831, ηp² = 0.001) or Standard waveforms (F(2, 261.4) = 0.16, p = 0.851, ηp² = 0.001). No main effects reached significance, indicating stable P1 amplitude across development for both conditions.

##### P1 Spatial Characteristics (Center of Mass)

The P1 component showed consistent posterior shifts in topographic distribution from T1 through T3. The total displacement was 0.078 units by T3, with the anterior-posterior component showing 0.054 units of posterior shift. This posterior displacement was accompanied by modest leftward (0.028 units) and inferior (0.048 units) shifts. The profile of spatial reorganization without a corresponding amplitude change suggests that the generators are reorganizing their geometry rather than showing a global increase or decrease in field strength. The posterior migration aligns with increasing contribution of specialized temporal auditory regions as they mature, and the leftward shift is consistent with evidence for early left hemisphere specialization for structured speech processing documented in neuroimaging studies.

#### 3.3.2. N1 Component

##### N1 Latency

For Deviant waveforms, a significant Session × Language interaction was observed (F(2, 244.6) = 3.12, p = 0.046, ηp² = 0.025), indicating differential developmental trajectories between native and nonnative conditions. This was the only significant interaction observed across all ERP measures. Significant main effects of Session (F(2, 276.4) = 11.59, p < 0.001, ηp² = 0.077) and Language (F(1, 244.8) = 5.23, p = 0.023, ηp² = 0.021) were also observed.

For Standard waveforms, the Session × Language interaction was not significant (F(2, 253.8) = 0.73, p = 0.482, ηp² = 0.006), although a significant main effect of Session emerged (F(2, 287.3) = 18.08, p < 0.001, ηp² = 0.112).

##### N1 Amplitude

No significant Session × Language interactions were observed for either Deviant (F(2, 260.4) = 1.23, p = 0.293, ηp² = 0.009) or Standard waveforms (F(2, 262.3) = 0.20, p = 0.820, ηp² = 0.002). No main effects reached significance.

##### N1 Spatial Characteristics (Center of Mass)

The N1 component exhibited a bidirectional pattern in topographic distribution for native consonants: posterior shift from T1 to T2 (0.039 units) followed by anterior shift from T2 to T3 (0.055 units), resulting in a net anterior displacement of 0.058 units from T1 to T3. For nonnative consonants, the N1 topographic distribution showed the largest spatial shift observed in the study, with total displacement of 0.099 units from T1 to T3. The trajectory included a small posterior shift from T1 to T2 (0.027 units) followed by a large anterior shift from T2 to T3 (0.089 units), demonstrating the same bidirectional pattern observed for native consonants but with greater magnitude. This broad spatial reorganization may reflect the competing effects of general auditory maturation and language-specific tuning on the cortical areas engaged for processing phonemic differences.

#### 3.3.3. P2 Component

##### P2 Latency

The Session × Language interaction was not significant for either Deviant (F(2, 237.3) = 1.61, p = 0.202, ηp² = 0.013) or Standard waveforms (F(2, 238.9) = 0.95, p = 0.390, ηp² = 0.008). A significant main effect of Session was observed for Standard waveforms (F(2, 272.0) = 6.51, p = 0.002, ηp² = 0.046). For Deviant waveforms, a significant main effect of Language emerged (F(1, 236.8) = 5.23, p = 0.023, ηp² = 0.022).

##### P2 Amplitude

No significant Session × Language interactions were observed for either Deviant (F(2, 251.3) = 0.57, p = 0.564, ηp² = 0.005) or Standard waveforms (F(2, 245.4) = 0.76, p = 0.471, ηp² = 0.006). Significant main effects of Session were observed for both waveforms (Deviant: F(2, 279.2) = 4.64, p = 0.010, ηp² = 0.032; Standard: F(2, 272.8) = 3.15, p = 0.044, ηp² = 0.023), indicating general maturational increases in P2 amplitude that did not differ by language condition.

#### 3.3.4. Summary of Spatial (Center of Mass) Results

Table 3 summarizes the center of mass displacement for each ERP component and condition across development. The P1 component showed consistent posterior and leftward migration, while N1 and MMR components exhibited anterior shifts, with nonnative consonants showing larger spatial reorganization for the N1 component.

**Table 3.** Center of Mass Displacement Summary.

| Peak | Condition | T1→T2 (units) | T2→T3 (units) | Total T1→T3 | Primary Direction |
| --- | --- | --- | --- | --- | --- |
| P1 | Combined | – | – | 0.078 | Posterior, Left |
| N1 | Native | 0.039 (post) | 0.055 (ant) | 0.058 | Net Anterior |
| N1 | Nonnative | 0.027 (post) | 0.089 (ant) | 0.099 | Net Anterior |
| MMR | Native | 0.018 (post) | – | 0.061 | Anterior |
| MMR | Nonnative | – | – | 0.065 | Anterior |

*Note:* Displacement values are in normalized units on a unit sphere. Directional labels: post = posterior, ant = anterior. Shifts $<0.02$ = minimal, $0.02-0.05$ = small, $0.05-0.10$ = moderate, $\geq 0.10$ = large.

### 3.4. Summary of Session × Language Interactions

Table 4 summarizes the Session × Language interaction tests across all ERP measures. Of the 14 interaction tests conducted (2 MMR measures + 12 peak measures), only one reached statistical significance: N1 latency for Deviant waveforms (p = 0.046).

**Table 4.**
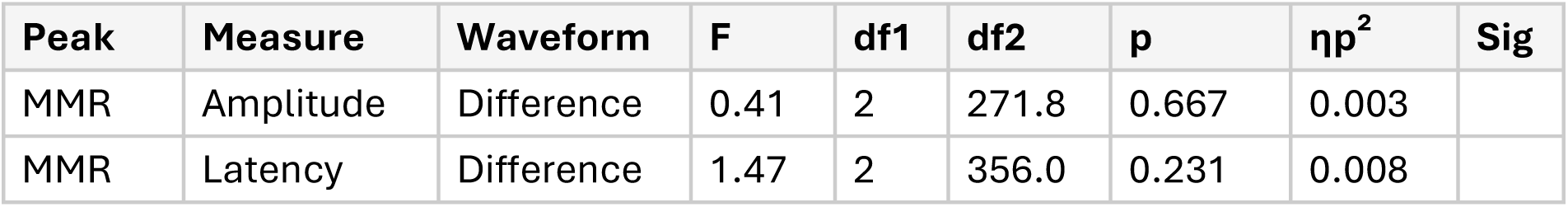
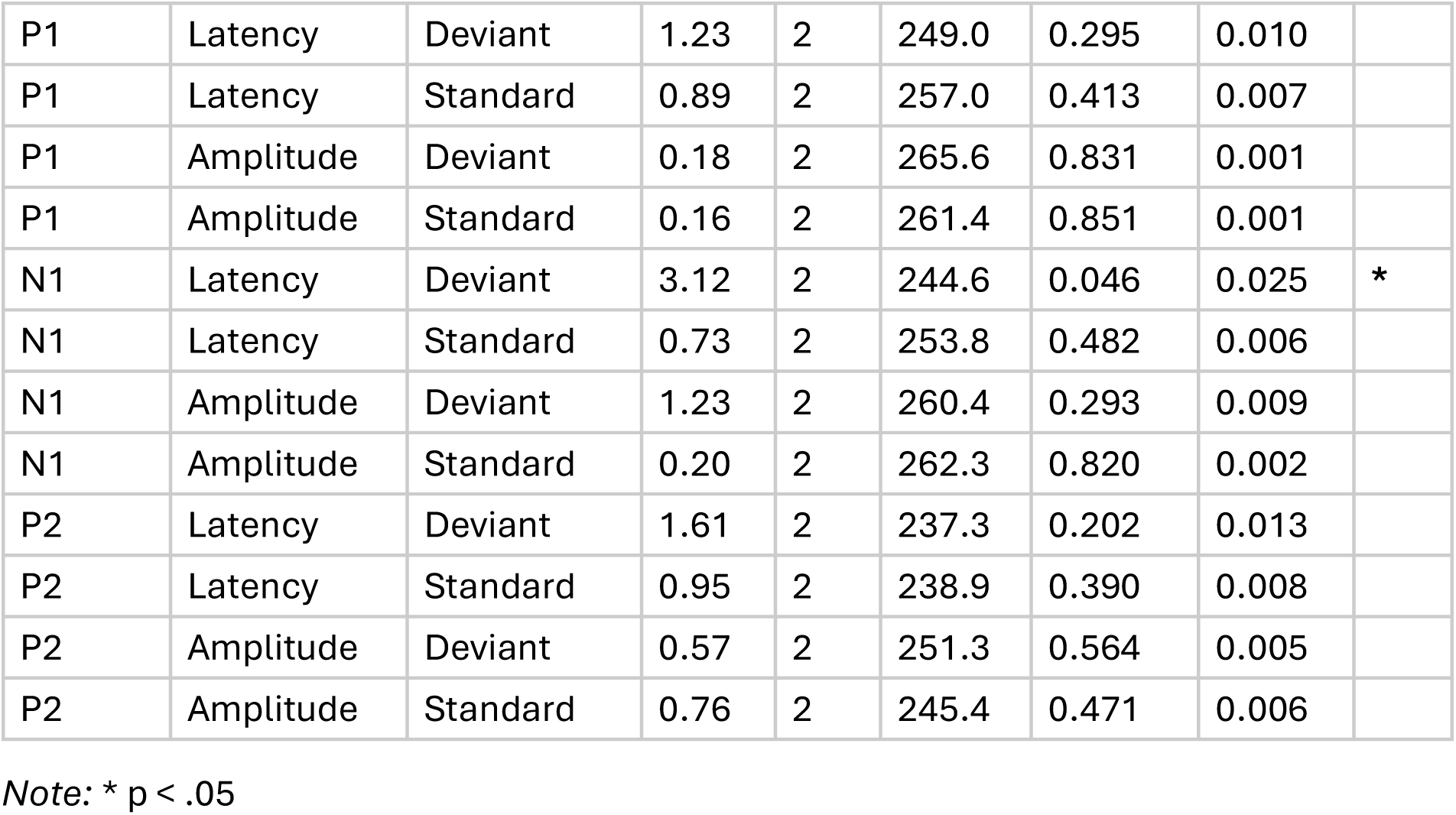
Summary of Session × Language Interaction Tests.

### 3.5. Bayes Factor Analysis

Bayesian analysis examined whether evidence strength for neural discrimination differed between native and nonnative conditions across development. The proportion of individual subjects showing strong evidence (BF > 10) for an MMR was uniformly low across all conditions (0% in all cells), precluding categorical analysis.

Analysis of log-transformed maximum Bayes Factors revealed a significant Session × Language interaction (F(2, 254.0) = 100.76, p < 0.001, ηp² = 0.442), indicating that the strength of evidence for neural discrimination followed different developmental trajectories for native and nonnative contrasts. Specifically, evidence strength increased from T1 to T2 for native contrasts but decreased for nonnative contrasts, with both conditions showing decreased evidence at T3.

## 4. Discussion

The present longitudinal study examined whether cortical speech processing develops differently for native versus nonnative consonant contrasts across the first year of life. Using ERPs collected at approximately 3, 6, and 12 months of age, we directly tested the prediction from perceptual attunement theory that neural responses to native speech should strengthen while responses to nonnative speech should weaken or remain unchanged over development.

Contrary to predictions from perceptual attunement theory, our primary finding was that most ERP measures showed similar developmental trajectories for native and nonnative consonant contrasts. Of the 14 Session × Language interaction tests conducted, only one reached statistical significance: N1 latency for deviant waveforms (p = 0.046, ηp² = 0.025). This pattern of results suggests that, at least for the consonant contrasts and age range examined, neural processing of native and nonnative speech sounds develops along largely parallel pathways.

### 4.1. MMR Development

The mismatch response, often considered the most direct neural index of discrimination, showed no differential development between native and nonnative contrasts. Neither MMR amplitude nor latency exhibited significant Session × Language interactions (both p > 0.2). This finding is inconsistent with studies reporting enhanced MMR to native contrasts and diminished MMR to nonnative contrasts during the first year (Cheour et al., 1998; Rivera-Gaxiola et al., 2005). However, it is consistent with more recent work suggesting that perceptual attunement may be more subtle and context-dependent than originally characterized (Maurer & Werker, 2014).

The lack of significant main effects of Session on MMR amplitude (p = 0.480) also indicates that neural discrimination abilities remained relatively stable across the 3-12 month period, rather than showing the strengthening trajectory often reported in the literature. This stability may reflect the relatively late emergence of robust consonant discrimination compared to vowel discrimination, or may indicate that our consonant contrasts (/ba/-/da/ and dental-retroflex) were both sufficiently salient that discrimination was already established by 3 months.

### 4.2. Obligatory ERP Components

The obligatory ERP components (P1, N1, P2) showed significant main effects of Session for several measures, indicating general maturational changes in auditory cortical processing. Specifically, P1 latency decreased significantly across development (both waveforms, p < 0.001), consistent with increasing efficiency of early sensory encoding. P2 amplitude increased significantly across development (both waveforms, p < 0.05), suggesting strengthening of later-stage auditory processing.

Critically, however, these maturational effects did not differ between native and nonnative conditions; the Session × Language interactions were uniformly non-significant (all p > 0.2) except for N1 latency in deviant waveforms. This pattern suggests that the observed maturational changes reflect domain-general auditory development rather than language-specific tuning.

Importantly, while P1 amplitude remained stable throughout development and no significant main effects of session were detected, the component’s center of mass was found to undergo systematic posterior migration (0.078 units by T3) and a leftward shift (0.028 units), which indicates hemispheric lateralization is starting to emerge. The profile of spatial reorganization without a corresponding amplitude change suggests that the generators are reorganizing their geometry rather than a global increase/decrease in field strength. The posterior migration aligns with increasing contribution of specialized temporal auditory regions as they appropriately mature, and the leftward shift is consistent with recent accumulation of evidence for early left hemisphere specialization on structured speech processing that has been documented in neuroimaging studies, even in very young infants (approximately three months of age). Behavioral research shows that early left hemisphere specialization can contribute to efficient auditory processing across different domains. Finally, the dissociation between stable amplitude and migrating generators suggests that the spatial location of the measurement may be as important as the magnitude of the response for characterizing the maturation of early auditory encoding.

### 4.3. N1 Latency

The sole significant Session × Language interaction was observed for N1 latency in deviant waveforms (p = 0.046, ηp² = 0.025). The N1 component has been linked to change detection and early attentional orienting to acoustic deviations. The finding that only deviant-evoked N1 latency showed differential trajectories is intriguing, as it suggests that language-specific effects may be most apparent in the timing of neural responses to unexpected phonetic changes rather than in overall response magnitude.

The small effect size (ηp² = 0.025) and borderline significance (p = 0.046) warrant cautious interpretation. Given the multiple comparisons conducted, this single significant result should be considered preliminary evidence requiring replication. Nevertheless, the specificity of this effect to deviant-evoked responses is theoretically meaningful, as perceptual attunement is thought to manifest primarily in how the auditory system responds to violations of expectation based on accumulated language experience.

### 4.4. Bayes Factor Analysis

While individual-level Bayes Factor analysis showed uniformly weak evidence for MMR (0% of subjects reaching BF > 10 in any condition), the analysis of evidence strength across subjects revealed a significant Session × Language interaction. This suggests that although robust individual-level discrimination was rare, the population-level pattern of evidence showed differential development. Native contrasts showed increasing evidence from T1 to T2, while nonnative contrasts showed decreasing evidence, consistent with perceptual attunement predictions.

This dissociation between the null findings for MMR amplitude/latency and the significant Bayes Factor interaction highlights the importance of examining multiple indices of neural discrimination. The Bayes Factor analysis, which captures the overall pattern of evidence across the entire MMR time window rather than just the peak, may be more sensitive to subtle shifts in discrimination processes.

### 4.5. Theoretical Implications

Our findings challenge the view that perceptual attunement produces robust, easily measurable changes in neural responses to native versus nonnative speech across the first year. Several interpretations warrant consideration:

First, perceptual attunement for consonants may occur later than the 12-month endpoint of our study. Behavioral research suggests that consonant discrimination undergoes substantial reorganization between 8 and 12 months (Werker & Tees, 1984), and neural signatures may lag behind behavioral changes or require additional time to consolidate.

Second, the magnitude of attunement effects may be smaller than previously reported, particularly when examined using within-subject designs that control for individual differences in overall ERP morphology. Cross-sectional studies comparing groups at different ages may overestimate attunement effects by confounding cohort differences with developmental change.

Third, perceptual attunement may be highly stimulus-specific, with stronger effects for contrasts that are more phonologically relevant or that show greater acoustic overlap with the infant’s primary language. The native contrast used in this study (/ba/-/da/) is acoustically robust and may be discriminated even without extensive language experience, limiting the potential for experience-dependent enhancement.

Fourth, it is possible that attunement to non-native contrasts may still occur in a natural environment, but that repeated test visits, listening to the same repeated phonemes, result in a phenomenon where the developing brain learns to discriminate the test sounds. In such a case, the brain treats these contrasts as behaviorally relevant, creating a situation that alters the normal course of attunement.

### 4.6. Limitations

Several limitations should be considered when interpreting these findings. First, the sample size at T3 (n = 46) was smaller than at earlier time points due to participant testing constraints, potentially limiting power to detect developmental effects at the 12-month time point. Second, the use of different EEG amplifiers across sessions, while normalized through preprocessing, introduces potential systematic variance. Third, the predominantly sleeping state of infants during recording may not fully capture the range of cortical processes engaged during active listening in natural environments. Finally, the specific consonant contrasts tested may not generalize to other phonetic distinctions that show stronger attunement effects.

## 5. Conclusion

The present longitudinal study examined developmental trajectories of cortical speech processing for native and nonnative consonant contrasts across the first year of life. Contrary to predictions from perceptual attunement theory, we found that most ERP measures, including MMR amplitude, MMR latency, and obligatory component measures, showed similar developmental trajectories for native and nonnative contrasts. Only N1 latency for deviant waveforms showed a significant Session × Language interaction, providing limited evidence for differential neural development.

These findings suggest that robust neural signatures of perceptual attunement for consonants may be more subtle, later-emerging, or stimulus-specific than commonly assumed. The significant main effects of Session on several ERP measures confirm ongoing auditory maturation during the first year, but this maturation appears to be domain-general rather than language-specific for the consonant contrasts examined. Notably, center of mass analysis revealed systematic spatial reorganization of ERP topography across development, with P1 showing posterior migration and leftward shifts suggestive of emerging hemispheric lateralization, even in the absence of amplitude changes. This dissociation between stable amplitude and migrating generators suggests that spatial location may be as important as response magnitude for characterizing auditory maturation. Future research should examine whether attunement effects are stronger for different contrasts, emerge at later ages, or require more sensitive analytical approaches to detect.

Practically, these findings suggest that equivalent neural responses to native and nonnative consonant contrasts during the first year should not necessarily be interpreted as evidence of atypical development. Rather, indices of general auditory maturation may be more reliable markers of healthy auditory development during early infancy.

## Acknowledgments

We would like to express our gratitude to all families, Colorado Home Intervention Program, and audiologists who participated in this project.

## Data availability statement

De-identified data will be made available after institutional review and approval.

Funding for this research was provided by the National Institutes of Health: National Institute on Deafness and other Communication Disorders K23DC01358, NIDCD and OBSSR R01-DC021068and by CCTSI=NIH/NCRR Colorado CTSI Grant Number UL1 TR001082 to author KU. National Institute on Disability, Independent Living, and Rehabilitation Research (NIDILRR #90RE5020) to author PMG; NIDILRR is a Center within the Administration for Community Living (ACL), Department of Health and Human Services (HHS), USA. The contents of this research manuscript do not necessarily represent the policy of NIDILRR, ACL, or HHS, and you should not assume endorsement by the Federal Government.

## Statement on the use of AI

All ideas, hypotheses, interpretations, and scientific conclusions are original work of the study authors. Generative AI tools, including Grammarly, MATLAB Copilot, and Claude (Anthropic), were used during manuscript preparation to assist with grammar, readability, and code documentation. All AI-generated suggestions were reviewed and edited by the authors, who take full responsibility for the final content.

